# Behaviorally irrelevant feature matching increases neural and behavioral working memory readout

**DOI:** 10.1101/2023.09.12.557327

**Authors:** Aytaç Karabay, Michael J. Wolff, Veera Ruuskanen, Elkan G. Akyürek

## Abstract

There is an ongoing debate about whether working memory (WM) maintenance relies on persistent activity and/or short-term synaptic plasticity. This is a challenging question, because neuroimaging techniques in cognitive neuroscience measure activity only. Recently, neural perturbation techniques have been developed to tackle this issue, such as visual impulse perturbation or “pinging”, which reveals (un)attended WM content during maintenance. There are contrasting explanations of how pinging reveals WM content, which is central to the debate. Pinging could reveal mnemonic representations by perturbing content-specific networks or by increasing the neural signal-to-noise ratio of active neural states. Here we tested the extent to which the neural impulse response is patterned by the WM network, by presenting two different impulse stimuli. If the impulse interacts with WM networks, the WM-specific impulse response should be enhanced by physical overlap between the initial memory item and the subsequent external perturbation stimulus. This prediction was tested in a working memory task by matching or mismatching task-irrelevant spatial frequencies between memory items and impulse stimuli, as well as probes. Matching probe spatial frequency with memory items resulted in faster behavioral response times and matching impulse spatial frequency with memory items increased the specificity of the neural impulse response as measured from EEG. Matching spatial frequencies did neither result in globally stronger neural responses nor in a larger decrease in trial-to-trial variability compared to mismatching spatial frequencies. The improved neural and behavioural readout of irrelevant feature matching provide evidence that impulse perturbation interacts directly with the memory representations.

---

Cognitive neuroscientists attempt to relate cognitive processes to neural activity that is highly complex and multidimensional. Compounding this challenge is the fact that neural activity is only part of the picture. Synaptic processes that alter the functional connectivity in the brain’s networks also play an important role. Such processes need not correlate strongly with neural firing, which makes them difficult to assess with non-invasive measurements. These difficulties have recently come to a head in the study of working memory (WM), where a debate has arisen about its neural basis (e.g., Masse et al., 2020; Sreenivasan et al., 2014).

Maintenance of information in WM was long thought to involve persistent neural firing, supported by evidence of ongoing neuronal spiking during memory maintenance in higher-order cortical areas, such as the prefrontal and parietal regions (Curtis & D’Esposito, 2003; Funahashi et al., 1989). However, recent studies suggested that this apparent “persistence” may be an artefact of trial averaging and temporal smoothing of short-lived oscillatory activity bursts interrupted by silent periods (Lundqvist et al., 2016; 2018; Miller et al., 2018). These silent periods may be bridged via changes in synaptic efficacy (Kozachkov et al., 2022; Mongillo et al., 2008; Stokes, 2015). Specifically, during the encoding stage, the neural response to the sensory input may leave behind a patterned “silent” neural trace supporting WM maintenance (Stokes, 2015). However, it has been argued these “silent” periods are null-findings, persistent neural activity during WM maintenance is still present, albeit ata level that is too low to be reliably picked by neuroimaging tools (Schneegans & Bays, 2017; Barbosa et al., 2021).

Importantly, regardless of the underlying neural mechanism of WM maintenance, probing these low-activity or synaptic neural states is challenging with conventional neuroimaging techniques, which all rely on robust activity-dependent signals. To overcome this issue, impulse perturbation (“pinging”) techniques were developed that have proven effective at illuminating WM contents during maintenance (Rose et al., 2016; Stokes et al., 2013; Wolff et al., 2015; 2017). The pinging method involves presenting an invariant, task-irrelevant impulse stimulus (or alternatively, a TMS pulse; Rose et al., 2016) during the WM delay, while a participant’s electroencephalogram (EEG) is recorded (Wolff et al., 2015; 2017). The impulse is thought to work analogously to sonar; as a surge of activation is driven through the network, the resulting pattern of activity reflects not only the invariant neural response to the impulse stimulus, but also the current state of the perturbed network. Given that the impulse response has been found to contain information about the contents of WM in various studies (e.g., Duncan et al., 2023; Kandemir & Akyürek, 2023; Wolff et al., 2015; 2017), this strongly suggests there is overlap between the network that responds to the impulse stimulus, and the network that contains WM-specific neural traces. Indeed, it has been found that while the neural impulse response to auditory stimulation during an auditory WM task contained information about the auditory WM content, using the same auditory stimulation in a visual WM task did not result in a WM-specific neural response (Wolff et al., 2020b). Similarly, decoding accuracy of unattended WM content was higher when TMS was targeted to content-specific brain regions compared to unrelated regions (Rose et al., 2016). This implies that the neural impulse response is WM-specific as long as the neural processing of the impulse stimulus interacts with the sensory-specific WM trace, which should be more likely when WM memoranda and impulse stimulus share sensory modalities or other features (in line with the idea of sensory recruitment for WM; Emrich et al., 2013; Pasternak & Greenlee, 2005; Postle, 2006).^1^

However, an alternative account has been put forward (Barbosa et al., 2021), which suggests that the primary effect of visual impulse perturbation is to improve the signal-to-noise ratio (SNR) (as measured from the trial-to-trial variability of the signal) of ongoing neural activity (but see Wolff et al., 2021). This may happen because the presentation of the impulse reduces the amount of variability in the EEG; sensory input is known to have a stabilizing effect on neural activity (Churchland et al., 2010; Pfurtscheller et al., 1979; 1994). If improving the SNR is indeed the only function of the impulse, it could mean that the impulse does not actually perturb the memory network, but more generically reveals items that are held in activity patterns hidden in noise.

Although the efficacy of the impulse perturbation technique is not questioned by this alternative account, it remains important to understand the neural mechanisms that produce its effects. Therefore, in our experiment, we aimed to assess to what degree the impulse response reflects a WM network-specific response (regardless of its activity-state), or alternatively, a global neural response. Two different visual impulse and probe stimuli were presented in a WM task. The memory items were orientation gratings, and participants were asked to judge whether their orientation was clockwise or counterclockwise (CW, CCW) relative to that of a probe stimulus, which was presented after a short delay at the end of each trial. Critically, the impulses and probes either matched or did not match the memory items on their spatial frequency, which was an integral^2^, but task-irrelevant feature (Fig 1a-b).

**Figure 1.**
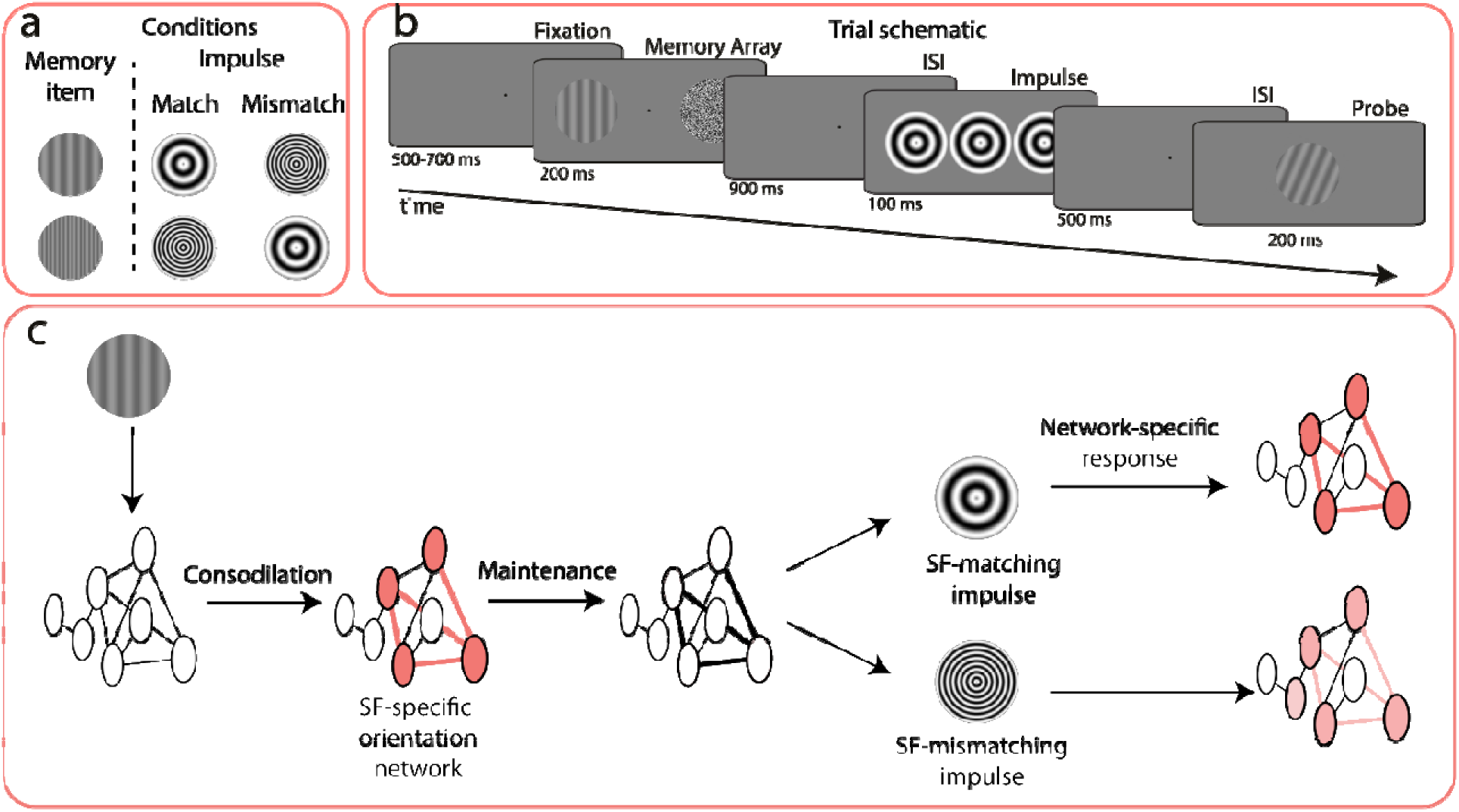
Illustration of the trial design and principal hypothesis. **a.** Experimental manipulation showing memory item-impulse spatial frequency (SF) match and mismatch conditions. **b.** Trial schematic. Participants were asked to gaze at the center of the screen. After a variable delay period, the memory array, which consisted of a low or high spatial frequency orientation grating and a noise patch (the minimum and maximum brightness of each pixel of the noise patch are exaggerated for illustration), were shown on either side of the fixation dot. The task was to remember the orientation, and judge whether it was clockwise or counterclockwise oriented relative to the probe. During the delay period, a visual impulse was shown. The spatial frequency of the impulse either matched or did not match the orientation grating. **c.** Hypothesis. A matching visual impulse perturbs WM content better due to its shared spatial frequency-specific neural code.

We hypothesized that if the impulse response reflects the perturbed WM network, as suggested by Wolff and colleagues (2017), a matching *impulse* should act like a matched filter and reveal the memory content with greater decoding accuracy because it targets the network storing the representation more precisely, regardless of whether WM content is maintained with persistent activity and/or STSP (Figure 1c). Similarly, a matching *probe* should improve behavioral readout by facilitating the transition towards the behaviorally relevant matched filter response (Myers et al., 2015; Stokes, 2015; Remington et al., 2018; Nikolic et al., 2009; Sugase-Miyamoto et al., 2008). The assumption that the same orientations may be represented within distinct neural traces in WM if their task-irrelevant features are different is supported by behavioral and neural findings, which suggest that objects are held in WM with all their features (Lin et al., 2021; Luck & Vogel, 1997), even when some are irrelevant (such as location: Zhou et al., 2022). As alluded to, this is furthermore in line with the sensory recruitment theory suggesting that WM representations are maintained in sensory regions that process the constituent features of the WM content (Emrich et al., 2013; Pasternak & Greenlee, 2005; Postle, 2006), and STSP-based accounts of WM, where any stimulation leaves behind an initial neural trace, whether relevant or not (Mongillo et al., 2008; Nikolic et al., 2009; Stokes, 2015).

Conversely, there should be no effects of impulse spatial frequency matching, if the previously reported WM-specificity of the impulse response is not due to a targeted perturbation of a WM-specific neural trace, but rather due to stimulus-induced, global neural noise reduction, which may generically facilitate the decodability of an active neural code (Barbosa et al., 2021; see also Churchland et al., 2010; Pfurtscheller et al., 1979; 1994), for example via phase reset (Wolff et al., 2021).

To preview the results, a matching impulse produced better decoding of mnemonic representations from the impulse-evoked multivariate EEG pattern than a mismatching impulse, and a matching probe reduced reaction times during memory recall. The stimulus-specific nature of the impulse response supports the idea that WM contents can be targeted directly by external perturbation (Stokes, 2015; Wolff et al., 2017). The positive behavioral effect of matching probes further suggests that the specificity of the impulse response is unlikely to be an epiphenomenon, but may rather reflect WM coding that is optimal for the task at hand (cf. Myers, 2022; Nairne, 2002). Finally, although the impulse also decreased trial-to-trial variability of the EEG voltage signal and decreased alpha power, these effects did not differ between match conditions, suggesting that neither of them can exclusively explain the impulse effect.

## Method

### Participants

A sample size of 30 was chosen following previous work using similar methodologies (Wolff et al., 2017; 2020a; 2020b). The sample size was not estimated prior to data collection. However, power calculations using G-power (Faul et al., 2007) was performed subsequently, with an alpha level of .05 and β of .80, following common guidelines (Lakens, 2013). These computations indicated that a sample size of 23 participants was required to detect medium to large effect sizes (Cohen’s *d*z = .614; based on decoding accuracy differences for cued items after auditory and visual stimulus presentations, Wolff et al., 2020b) in our T-tests (the main analysis of the manuscript). Our sample size aligns with these calculations, ensuring that the observed effects are not due to a lack of statistical power. In total, 31 participants, most of whom were undergraduate students, participated at the University of Groningen in exchange for monetary compensation (8 euros per hour). One participant was excluded due to excessive EEG noise (more than 40% of the trials were contaminated). In the final sample, there were thus 30 participants (18 females [16 right-handed], 12 males [12 right-handed], mean age = 23.3, range = 16-32). All participants reported normal or corrected-to-normal vision and signed an informed consent form prior to participation. The study was conducted in accordance with the Declaration of Helsinki (2008).

### Apparatus and stimuli

Participants were individually seated in dimly lit sound-attenuated testing cabins, approximately 60 cm from a 27-inch Asus IPS monitor (model VG279QM). The resolution was set to 1920 by 1080 pixels, at 16-bit color depth with a 100 Hz refresh rate. The monitor was not gamma-corrected or calibrated for luminance linearity, which may result in minor deviations in luminance contrast. OpenSesame 3.2.8 (Mathôt et al., 2012) with the Psychopy back-end (Pierce et al., 2019) was used for trial preparation and data collection, running under the Microsoft Windows 10 operating system. Responses were collected with a standard keyboard.

The stimuli were presented on a grey background (RGB = 128, 128, 128). A black fixation point (RGB = 0, 0, 0) was maintained in the center of the screen throughout the trial. Memory items and probes were gray sine-wave gratings presented at 20% contrast calculated by the Michelson Contrast, with a diameter of 5.93° of visual angle (Figure 2a). Spatial frequencies of the memory items, probes, and impulse stimuli were either 0.5 cycles/degree (low spatial frequency) or 1.4 cycles/degree (high spatial frequency), depending on the experimental condition (Figure 2b). Memory items (orientation gratings) were presented on the left or right side of the fixation dot, at an eccentricity of 5.93° of visual angle. A noise patch was shown at the same location on the other side of the fixation dot, matching the contrast and overall luminance of the memory items. Orientations were randomly chosen from 144 possible unique angles separated evenly between 0° and 179° with 1.25° separation. The angle difference between the memory item and the probe was chosen from 12 possible unique values, ranging from −40° to 40° (±5°, ± 10°, ± 16°, ± 24°, ± 32°, ± 40°, as in Wolff et al., 2017). Impulse stimuli consisted of three adjacent bull’s eyes presented at the center of the screen. Each bull’s eye was the same size as the memory items and probes (i.e., a diameter of 5.93° of visual angle).

**Figure 2.**
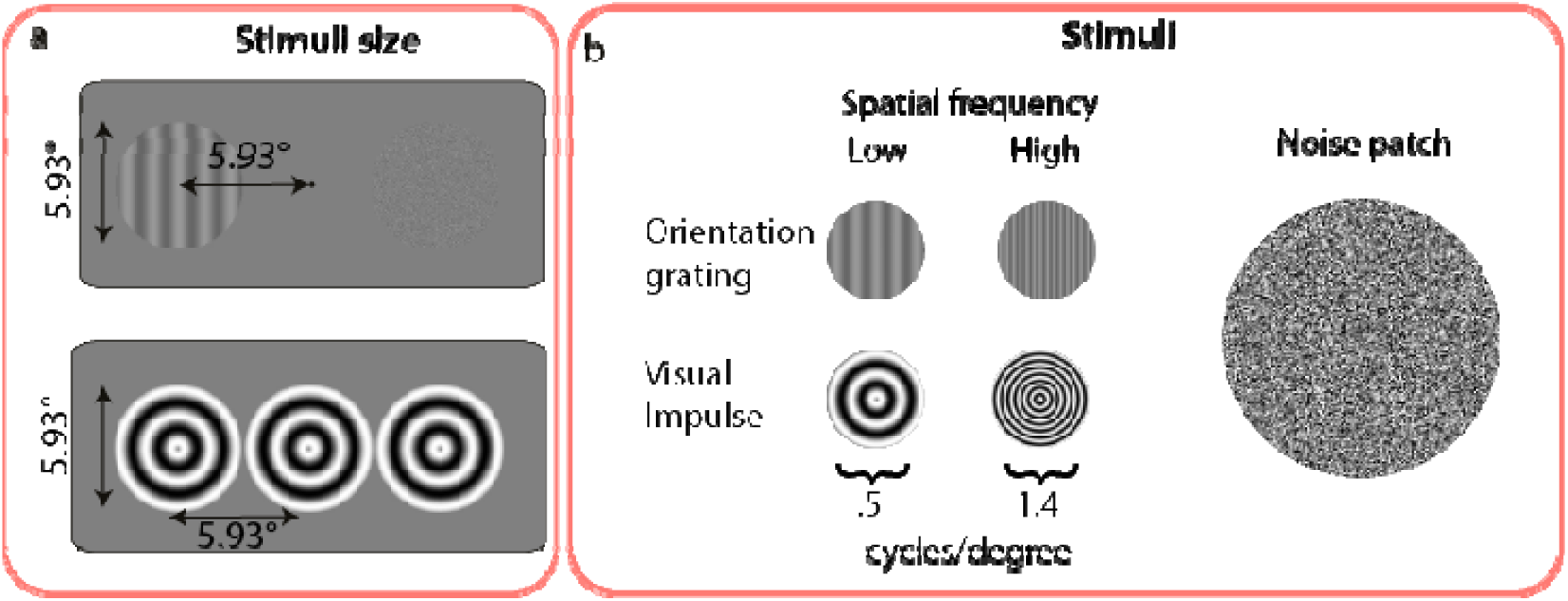
Stimulus details in degrees of visual angle (°). **a**. Size and location of low and high spatial frequency memory items and impulses. **b.** Spatial frequencies of memory items and impulses, and noise patch. For illustration purposes, the minimum and maximum brightness of each pixel of the noise patch are exaggerated.

### Procedure

Participants completed a visual WM task (Figure 1b) while EEG was recorded. The task was to report whether the probe was rotated clockwise or counterclockwise relative to the memory item. Each trial began with a fixation cross with random temporal jitter, such that it was presented for 500-700 ms. Next, the memory item and noise patch were simultaneously presented for 200 ms. After a 900 ms delay, the impulse stimulus was shown for 100 ms. After a 500 ms ISI, the memory probe was presented for 200 ms, and participants were required to give a speeded response by pressing m or c (counter-balanced between participants) within 1000 ms.

Trial-wise feedback was provided for 200 ms (a happy smiley for the correct response and an unhappy smiley for the incorrect response). There was a 500 ms inter-stimulus interval between trials. Participants started the experiment with 64 practice trials. If they failed to perform above 60% accuracy, they were asked to redo the practice trials of the experiment. Participants were given a chance to have a break between each block and each session (see below). After each block, the average accuracy that the participant had achieved within was shown. The total duration of the study was four hours, including EEG capping.

### Design

The principal variables of interest were whether the spatial frequency of the memory item matched that of the impulse and/or that of the probe (Figure 1c). The design was nevertheless also randomized and counterbalanced for the location of the memory item (left or right), the spatial frequency of the memory item (low or high), the spatial frequency of the impulse stimulus (low or high), and the spatial frequency of the probe stimulus (low or high). For each of the 16 resulting design cells, 144 unique orientations were shown. In addition, each probe difference was randomly assigned to those 144 unique orientations, which was counterbalanced within and between cells. Therefore, there were 2304 trials in total (16 * 144), which were separated into four consecutive sessions. Each session (576 trials) consisted of 24 blocks, and each block consisted of 24 trials. After randomization for these 2304 trials was done for each participant, the trial order was shuffled and separated into four sessions. The randomization of stimuli and conditions was achieved with experimental code written in R 4.0 (R Core Team, 2020).

### Behavioral data analysis

Mean accuracy was calculated for each participant across all trials for each condition of interest. To visualize the proportion of clockwise responses, counterclockwise responses were reverse-signed when the probe was counterclockwise relative to the memory item. Reaction time was calculated by retrieving median values for each subject per condition, but trials in which no response, a response that was too fast (<150 ms), or an incorrect response were given were discarded from the analysis.

### EEG acquisition

The EEG signal was acquired from 64 Ag/AgCl electrodes and one external EOG electrode using an equidistant hexagonal layout. The ground electrode was placed on the participants’ upper back. Electrodes placed above and below the left eye (EOG and 1L) and the temples (1LD and 1RD) were used for bipolar electrooculography. The impedance of all. The signal was recorded at 1000 Hz using an ANT Neuro Ω Eego Mylab amplifier and the associated software. During recording, the EEG was referenced to the electrode 5Z, and an antialiasing filter with a power cutoff at 260L

### EEG preprocessing

Using EEGLAB 2021 (Delorme & Makeig, 2004), the EEG signal was re-referenced to the common average of all electrodes and downsampled to 500 Hz. A high-pass filter of 0.1 Hz and a low-pass filter of 40 Hz were applied to the EEG signal. Noisy EEG channels were interpolated with the *pop_interp* function of EEGLAB. As a result, in four participants’ one to five EEG channels were replaced through spherical interpolation. Data were epoched relative to impulse onset (−1550 ms to 2000 ms). Following that, independent component analysis was run to remove eye-movement-related (blinks and saccades) components from the EEG signal. Trials with other EEG artefacts were detected semi-automatically using the *ft_rejectvisual* function of Fieldtrip 2017 (Oostenveld et al., 2011), and excluded from all subsequent analyses.

Wolff and colleagues’ (2020b) methodology was used for preprocessing EEG data for distance-based multivariate pattern analysis (MVPA). For the time-course decoding analysis, a sliding window approach was used, since pooling information over time may improve decoding accuracy (Grootswagers et al., 2017; Nemrodov et al., 2018). EEG data was downsampled to 100 Hz by taking the average of every 10 ms. Next, the data for each channel were normalized by subtracting the average voltage values within a sliding 100 ms time window from each voltage value (prior to each value), to remove stable activity that does not change within the time window. This normalization process emphasizes evoked dynamics by removing low-frequency, sustained activity. This is critical to our analysis as we are specifically interested in the transient, stimulus-evoked responses rather than ongoing background activity (Wolff et al., 2020a; 2020b). Therefore, only evoked dynamics were considered in this analysis. The 10 time-points within each time-window were combined for each channel and used as separate dimensions, resulting in 610 dimensions in total (61 channels (excluding the bipolar EOG channels) by 10 time-points). The sliding time-window was centered at 50 ms, such that a given time-point included information from −50 ms to +50 ms relative to it.

As a secondary analysis, information over the entire pre-selected time window of interest was pooled. Following Wolff et al. (2020a; 2020b) the time window of interest was determined as 100 to 400 ms relative to the onset of stimulation (memory item, impulse) a priori. After downsampling the EEG data to 100 Hz, data was normalized by subtracting average voltage values within the time window of interest for the same reason as described above. This resulted in 30 values per channel, and each value was considered as a separate dimension (61 channels, excluding the bipolar EOG channels, by 30 values, 1830 in total), which were used in the decoding analysis, resulting in a single decoding accuracy value per participant.

### Orientation decoding

Following Wolff and colleagues (2020a; 2020b), Mahalanobis-distance-based MVPA (De Maesschalck et al., 2000) was applied to decode potentially parametric memory representations of memory item orientations, where it is assumed that more dissimilar orientations result in more dissimilar EEG voltage patterns. Although memory orientations were counterbalanced, given that a different number of trials were discarded from the EEG analysis due to artefacts, an 8-fold procedure with subsampling to equalize unbalanced orientation distributions between conditions was used. This subsampling approach prevents the classifier from becoming biased toward conditions with more trials, ensuring that the overrepresentation of any condition is mitigated. Trials were assigned to the closest of 16 evenly split orientation bins (variable, see below) and randomly split into eight folds, seven of which constituted the train set, and one of them the test set. The number of trials in each orientation bin in each train set was equalized by random subsampling, matching the minimum number of trials in any orientation bin to ensure an unbiased training set. Next, the covariance matrix was computed with the train sets (seven folds), using a shrinkage estimator (Ledoit & Wolf, 2003) to capture the relationship of neural representations of orientation bins, while improving stability and accuracy in high-dimensional data with limited trials. The activity of each orientation bin in the subsampled train trials was averaged to generate a representative neural pattern for each bin by reducing trial-by-trial noise. The average bins of the train trials were convolved with a half cosine basis set raised to the 15^th^ power in order to pool information across similar orientations (Myers et al., 2015). Finally, to assess how closely each test trial matched the orientation-specific neural pattern, Mahalanobis distances of each trial in the test fold were computed relative to the averaged and basis-weighted orientation bins. This resulted in 16 distances per test trial, which were mean-centered per trial to remove any bias in distance magnitude across trials and ensure that the decoding reflected relative differences between conditions rather than absolute values. This procedure was repeated for all test and train fold combinations.

Because trials were subsampled, this procedure was repeated 100 times for each of eight different orientation spaces (0° to 168.75°, 1.40625° to 170.1563°, 2.8125° to 171.5625°, 4.2188° to 172.9688°, 5.625° to 174.375°, 7.0313° to 175.7813°, 8.4375° to 177.1875°, 9.8438° to 178.5938°, each in steps of 11.25°) to minimize any bias from fixed binning. For each trial, 800 samples for each of the 16 Mahalanobis distances was obtained. Distances were averaged per trial, ordered with regard to the orientation difference, and sign-reversed, such that larger values reflect larger pattern-similarity between the test trial and the averaged orientation bin. The pattern similarity curve was summarized into a single decoding value by taking the cosine vector mean, such that a higher value reflects higher pattern similarity between similar orientations and lower similarity between dissimilar orientations. This method effectively handles the circular nature of orientation data by measuring the angular differences between orientations and neural patterns, ensuring that orientations close to 0° and 180° are properly accounted for. By averaging the cosine similarities, this approach provides a single metric that captures how well the neural activity aligns with the stimulus orientations across all trials.

### Spatial frequency decoding

The spatial EEG activity pattern between 100 to 400 ms after memory item and impulse onset was used in spatial frequency reconstruction. The approach was similar to that of orientation reconstruction. Since there were only two spatial frequency conditions, the Mahalanobis distance of the spatial frequency condition relative to both spatial frequency conditions was computed. Following that, the Mahalanobis distance of the same spatial frequency condition was subtracted from that of the different spatial frequency condition. Positive values were converted to hits, and negative values were converted to misses. Hits and misses were averaged and presented as decoding accuracy in %.

### Variance reduction analysis

EEG data were epoched from −200 to 500 ms relative to impulse onset. Trials were divided into two groups with regard to memory item – impulse spatial frequency pair conditions. Next, each EEG channel was averaged for each condition per participant, and the variance of each channel’s activity was calculated on each time point of the EEG data. The variance of all channels was averaged for each time point. Following that, the relative change of variance was computed using the following formula:

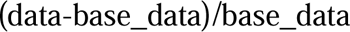

Where base_data was the variance during the baseline period (−200 to 0 ms relative to impulse onset) and data was the variance of the whole epoch.

The variance difference between match and mismatch conditions was calculated by subtracting one condition from another. A cluster-corrected permutation was run on the variance difference between conditions. Further, the average variance was estimated between 100 – 400 ms relative to impulse onset, since this time window was used for decoding memoranda. One extra participant was discarded from this analysis because the average variance in one condition was 2.5 standard deviations greater than that of the whole sample.

### Alpha power reduction analysis

Alpha power reduction between matching and mismatching impulses was compared. Alpha power was first computed by filtering the voltage data of all EEG channels between 8 and 12 Hz and Hilbert transforming the result using the following code in Matlab:

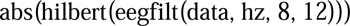

The filter length was 375 ms. The filtered data was subsequently averaged across all channels, before computing relative alpha power change after impulse presentation, using the same formula as for relative variance change ((data-base_data)/base_data)). Baseline alpha power was taken from the filtered data from −500 to −200 ms relative to impulse onset.

### Event-related potentials

To assess whether there was a univariate change in the visual sensory evoked potential between impulse conditions, the electrodes closest to the classic PO7 & PO8 scalp locations (5LC & 5RC) were examined. EEG was epoched from −200 to 500 ms relative to impulse onset, followed by a baseline correction from −200 ms to 0 ms (presentation of impulse). Trial epochs were then averaged per subject and condition, per time point. The ERP difference between match and mismatch conditions was calculated by subtracting one condition from another.

### Significance testing

A permutation T-test with 9999 Monte Carlo permutations was run for the analysis of behavioral data (response time and accuracy) using R (R Core Team., 2020), and decoding accuracy of pooled data using MATLAB. R was used to preprocess behavioral data and ggplot2 (Wickham, 2016) to plot outcomes. A cluster-corrected, one-sample permutation T-test with 100000 permutations was used for all time-course analyses. All tests were two-sided.

## Results and discussion

### Behavioral performance

As expected, accuracy increased, and RT decreased, as a function of the absolute difference between the orientation of the memory item and probe (Figure 3a-b). Average accuracy was 77.4%, with a standard deviation of 7.6%. Mean of medians probe RT was 536.1 ms, with a standard deviation of 100.1 ms.

**Figure 3.**
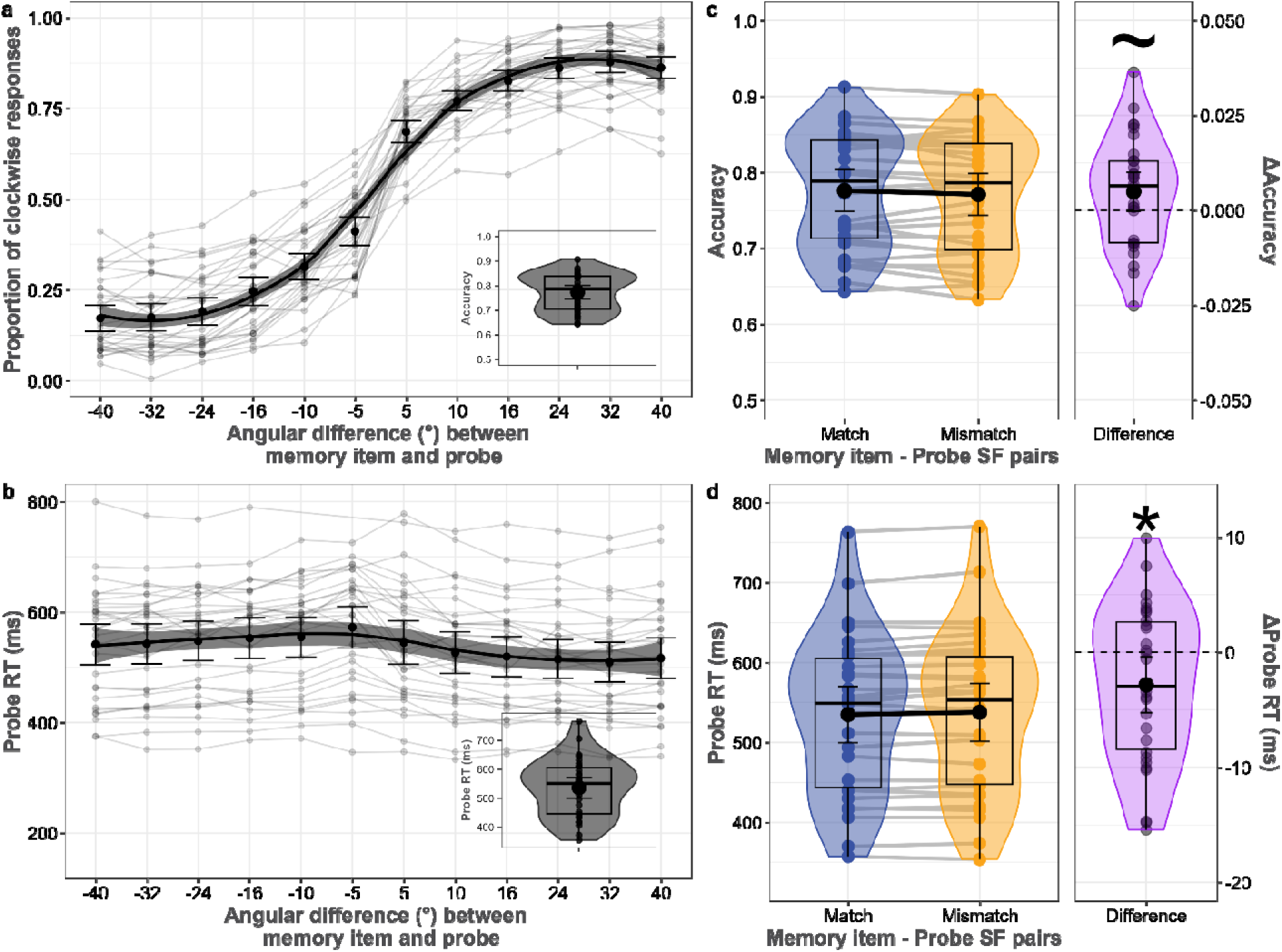
Behavioral performance. **a**. The proportion of clockwise responses and **b.** probe response time as a function of the angular difference between the memory item and probe. The inset plot on the top panel shows average accuracy, and the bottom panel shows average probe response time. **c.** Mean accuracy of memory item and probe spatial frequency pairs. **d.** Probe response time of memory item and probe spatial frequency pairs. Colored and grey dots represent individual data, black dots reflect averages, and error bars represent 95% confidence intervals calculated as 1.96 times the standard error of the mean. Transparent lines match subjects. Whisker plots show median and quartile values, and violin plots show the distribution. ∼ shows p < .1, * shows p < .05, ** shows p < .01, and *** shows p < .001 (permutation tests).

A marginally significant difference in accuracy existed between spatial frequency pairs of memory item and probe (*p* = .07; Figure 3c). Accuracy was 77.6% in the match (*sd =* 7.6%), and 77.1% in the mismatch (*sd =* 7.7%) condition. Participants responded faster when the memory item’s spatial frequency matched with the probe (*p* = .028; Figure 3d). Response time was 534.9 ms (*sd =* 99 ms) in the match condition, and 537.7 ms in the mismatch condition (*sd =* 95.7 ms). This provides behavioral evidence that matching a feature between memorandum and probe stimulus facilitates WM readout, even when the matching feature dimension is behaviorally irrelevant.

### Decoding memory item orientation in match and mismatch conditions

Cluster-corrected permutation tests showed significant clusters of memory item orientation decoding during the encoding period, in both the match (76 – 488 ms, *p* < .0001; Figure 4a-b left panel) and mismatch conditions (56 – 548 ms, *p* < .0001). Confirming the obvious fact that the memory item - impulse spatial frequency pairs can only influence decoding accuracies after impulse onset, there were no differences between conditions at any time point before impulse onset (*p* > .65 at all time points). The analysis on the whole time window of interest (100 to 400 ms) showed significant decoding for both the match and mismatch condition (*p* < .0001; *p* < .0001, respectively; Figure 4c top panel), with no difference between them (*p* = .216).

**Figure 4.**
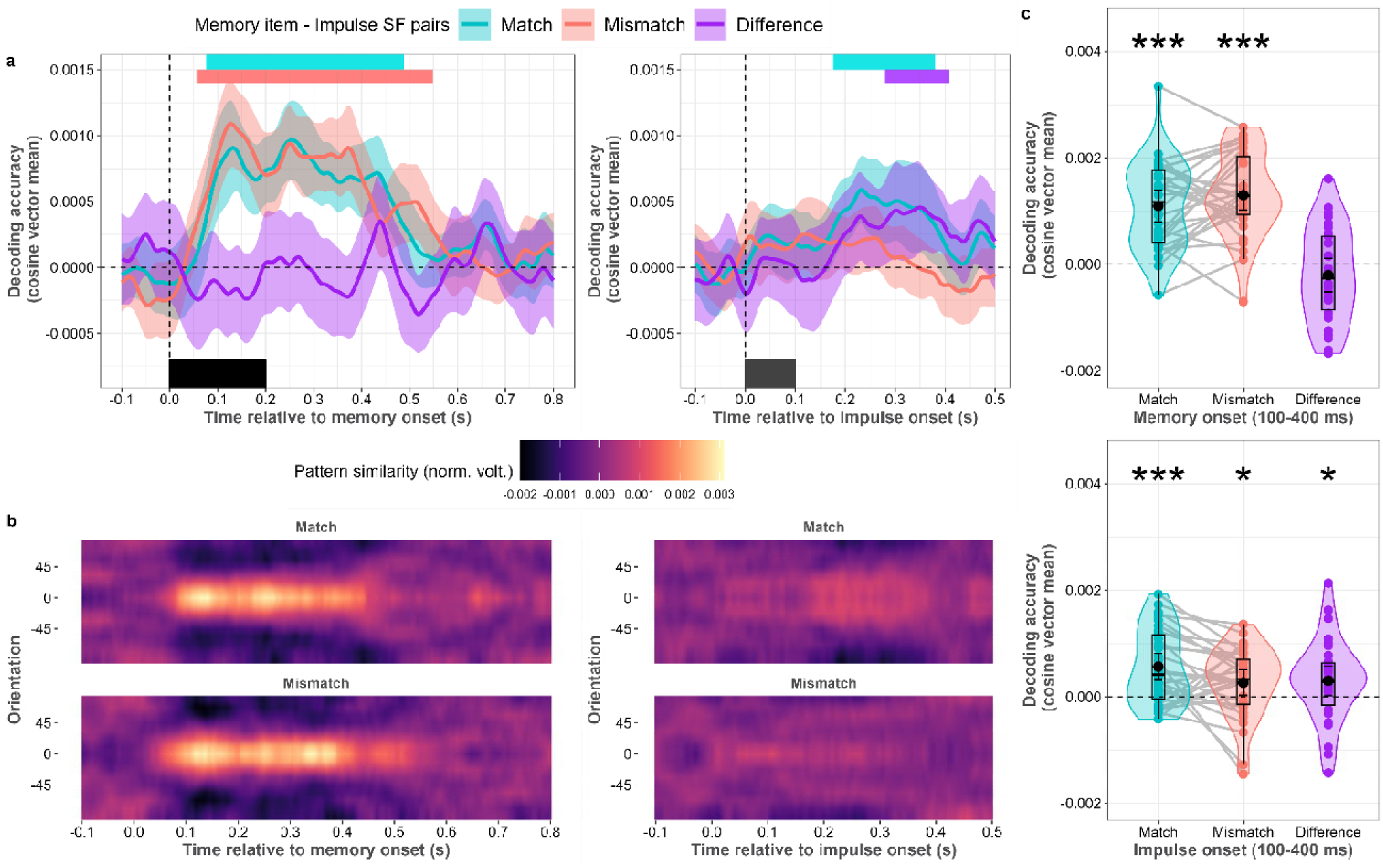
Orientation decoding of memory item – impulse spatial frequency (SF) pairs, during encoding and maintenance. **a.** Normalized average pattern similarity (mean-centered, sign-reversed Mahalanobis distance) of the neural dynamics at each time point for both conditions during encoding (post-memory item presentation; left plot), and during maintenance (post-impulse presentation; right plot). Colored horizontal bars indicate significant clusters of orientation decoding. The black rectangles show the onset and offset of the memory item or impulse stimulus **b.** Average distance to template of all angle bins (mean-centered, sign-reversed Mahalanobis distance) of the neural dynamics at each time point for each condition after memory item presentation, and impulse presentation. **c.** Orientation decoding relative to memory onset, and impulse onset in the time window of interest (100 – 400 ms relative to onset). Boxplots, violin plots, dots, lines, error bars, and asterisks follow Figure 3.

Time course decoding after impulse onset showed a significant cluster for the match condition (154 – 374 ms, *p* = .001; Figure 4a-b right panel), as well as a significant difference between match and mismatch conditions (278 – 398 ms, *p* = .01). The mismatch condition failed to reach significance. The spatio-temporal analysis on the whole time window of interest (100 to 400 ms) nevertheless showed significant decoding for both the match and mismatch condition (*p* < .001; *p* = .038, respectively; Figure 4c bottom panel), as well as a significant difference between them (*p* = .04).

The observed impulse-matching effect could be interpreted as sensory-driven, as impulses may perturb neural pathways related to sensory processing rather than WM. In such a case, the matching-dependent impulse response might reflect changes in synaptic efficacy within sensory pathways, driven by feature-specific sensory processing in neurons tuned to SF-specific orientations (Daugman, 1980; Webster & de Valois, 1995). Nonetheless, we conducted further control analyses to investigate the sensory account further (see Appendix for details). First, if the above-chance decoding accuracy after impulse presentation is driven by sensory processing, the decoding accuracy of memory orientations during the encoding period should influence the decoding accuracy during maintenance, as the decoder would be trained and tested on the same/similar sensory-driven neural processing. In this case, the correlations would be greater in the match condition as matching impulses would better perturb sensory-driven neural processes than WM-related neural processing due to the overlap of SF between matching impulses and memory orientations. However, this was not the case, as the difference was not significant, and the correlations were very low (see Appendix Figure A1). Second, if sensory-driven neural signals could explain the observed matching effect, neural signals would be expected to cross-generalize better in the match condition compared to the mismatch condition. However, this was also not the case, as there was no significant difference between the match and mismatch conditions, and none of the cross-generalizations reached significance (Appendix Figure A1b). We should note that the absence of cross-generalization could still result from differences between purely sensory-driven neural processing and the interaction of these processes with impulse presentations. Finally, we reanalyzed the Experiment 1 dataset from Wolff et al. (2017). In this experiment, the use of a retro-cue made sure that impulse-based decoding could be ascribed unambiguously to working memory, and indeed the paper found no impulse response for the uncued item. However, the original analysis was not based on the present dynamic baselining procedure, which isolates dynamic EEG components, and which might introduce sensory factors into the impulse response. We therefore applied the present decoding analysis to both cued (to-be-remembered) and uncued (to-be-ignored) memory items, and replicated the finding that only the to-be-remembered orientations were decodable (Appendix Figure A1c), precluding a sensory account of the present findings. Taken together, the results of these analyses do not support the idea that the current neural impulse response is related to sensory history.

### Testing for potential multivariate and univariate differences between memory item– impulse spatial frequency pairs

We were interested if there were any global differences between matching and mismatching impulse stimuli that might be related to the increased decoding of matching impulse trials. To this end, several univariate measures of known stimulus-evoked neural responses between matching and mismatching impulses, such as event-related potentials, across-trial variance reduction, and alpha power reduction, which are all related to one another, were compared (e.g., Arazi et al. 2017; Wolff et al. 2021).

There were no statistically significant differences in event-related potentials as measured from electrodes closest to the conventional PO7 and PO8 scalp locations (5LC and 5RC; *p* = 1, Figure 5a). There were also no statistical differences in relative variance reduction (*p* = 1 at all time points, Figure 5b), nor in relative alpha power reduction (*p* > .163 at all time points, Figure 5c). Impulse presentation reduced variance and alpha power in both match and mismatch conditions after impulse offset. Relative variance change after impulse onset was significant between 2 – 124 ms and 222 – 476 ms in the match condition (*p* = .046 and *p* = .007, respectively; Figure 4b left panel), and between 94 – 500 ms in the mismatch condition (*p* = .042 and *p* = .009, respectively; Figure 5b left panel). Relative alpha power change after impulse onset was significant between 224 – 500 ms in the match condition, and between 226 – 500 ms in the mismatch condition (*p* < .0001, *p* < .0001, respectively; Figure 5c left panel).

**Figure 5.**
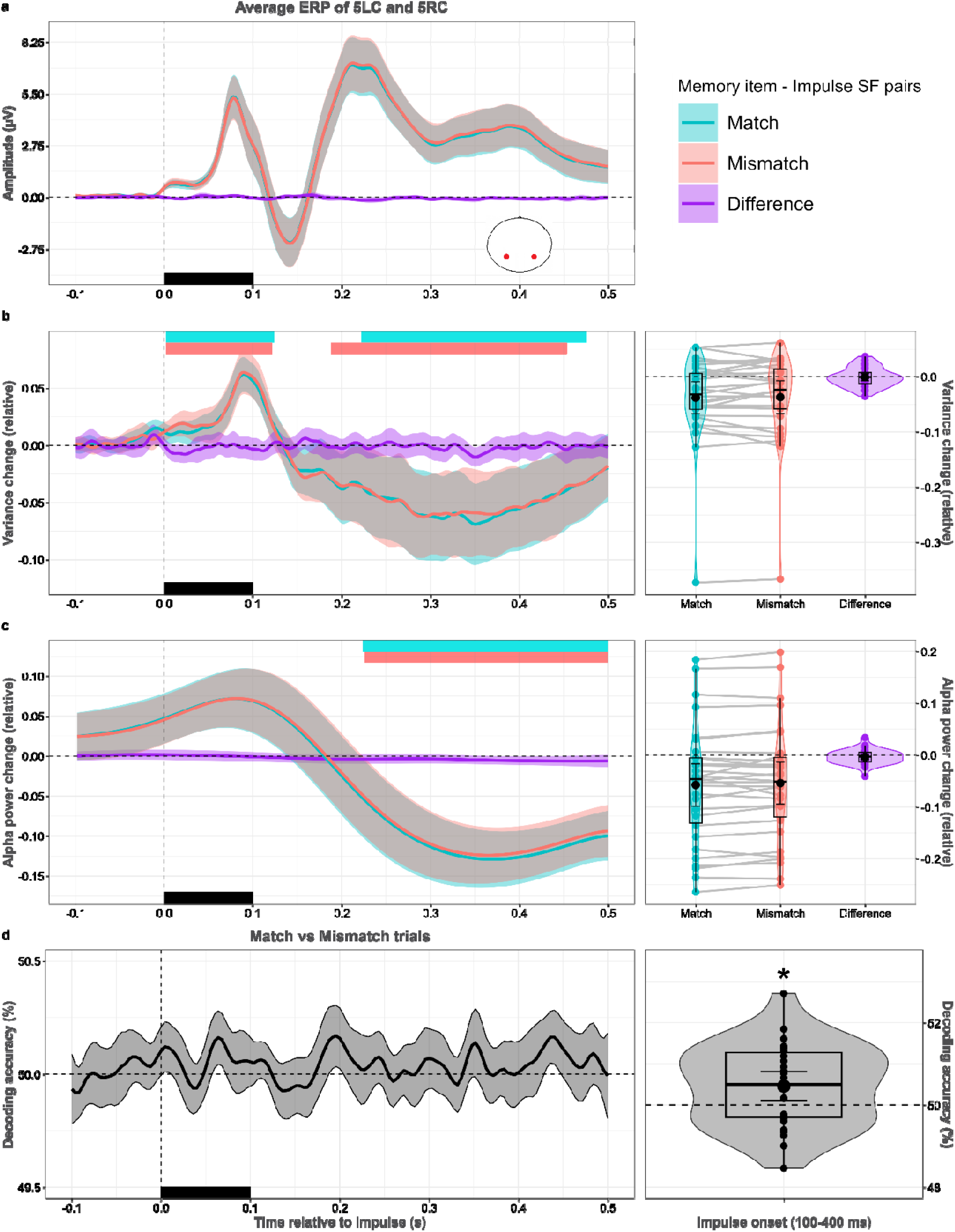
Univariate differences and condition decoding of match and mismatch trials. **a.** Impulse-locked event-related potentials in match and mismatch conditions. **b.** Left panel shows relative EEG variance for each time point. Right panel shows relative EEG variance change in the time window of interest (100 – 400 ms after impulse onset). **c.** Left panel shows relative alpha power change for each time point after impulse onset. Right panel shows relative alpha power change in the time window of interest. **d.** Left panel shows the decoding accuracy of experimental conditions (match vs. mismatch trials) per time point. Right panel shows the decoding accuracy of experimental conditions in the time window of interest. Colored rectangles on the top side of the left panels show significant clusters. Black rectangles show the onset and offset of the impulse. Colors, error bars, box plots, violin plots, dots, asterisks and lines follow Figure 3.

Finally, the question of whether there is a more subtle, multivariate difference between matching and mismatching impulse trials was investigated. While the time-course decoding showed no significant match/mismatch decoding (*p* > .40 at all time points, Figure 5d left panel), the time window of interest decoder did reach significance (*p* = .016, Figure 5d right panel). This suggests that there are no global differences between matching and mismatching impulse stimuli that could explain the WM-content decoding difference reported above. There was nevertheless a subtle multivariate difference between matching conditions, suggesting that there may be unique neural signatures or patterns that are activated in the presence of matching versus mismatching impulses, without changing the global response amplitude.

### Decoding memory item and impulse spatial frequency during encoding and maintenance

We furthermore tested whether the task-irrelevant spatial frequency of the memory item can be decoded during WM encoding and maintenance. The memory item’s spatial frequency was significantly decodable during WM encoding (*p* < .0001; Figure 6a). During WM maintenance, decoding of both the spatial frequency of the memory item and that of the impulse was also significant (*p* = .001, *p* < .0001, respectively; Figure 6b).

**Figure 6.**
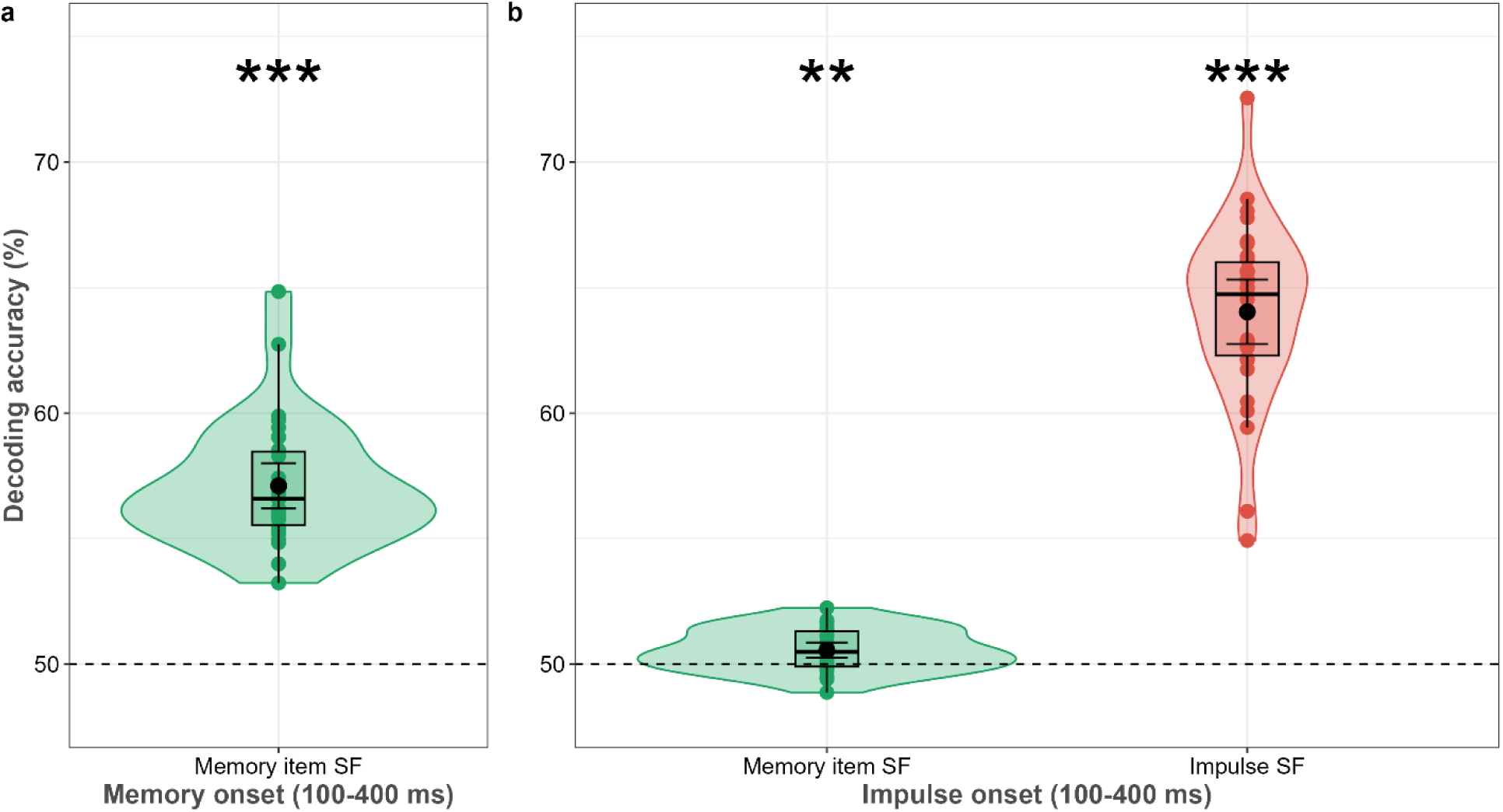
Decoding accuracy (%) of the memory item and impulse spatial frequency (SF) during. **a.** encoding and **b.** maintenance. Boxplots, violin plots, dots, error bars, and asterisks follow Figure 3.

### Decoding the location of the memory item during encoding and maintenance

Our final analysis tested whether the task-irrelevant spatial location of the memory item can be decoded during WM encoding and maintenance. Permutation T-tests showed that the location of the memory item was significantly decodable during WM encoding and maintenance (*p*s < .001; Figure 7). This finding was consistent with the decoding of SF, and suggests that all features of the memory items, task-irrelevant or not, were maintained in WM.

**Figure 7.**
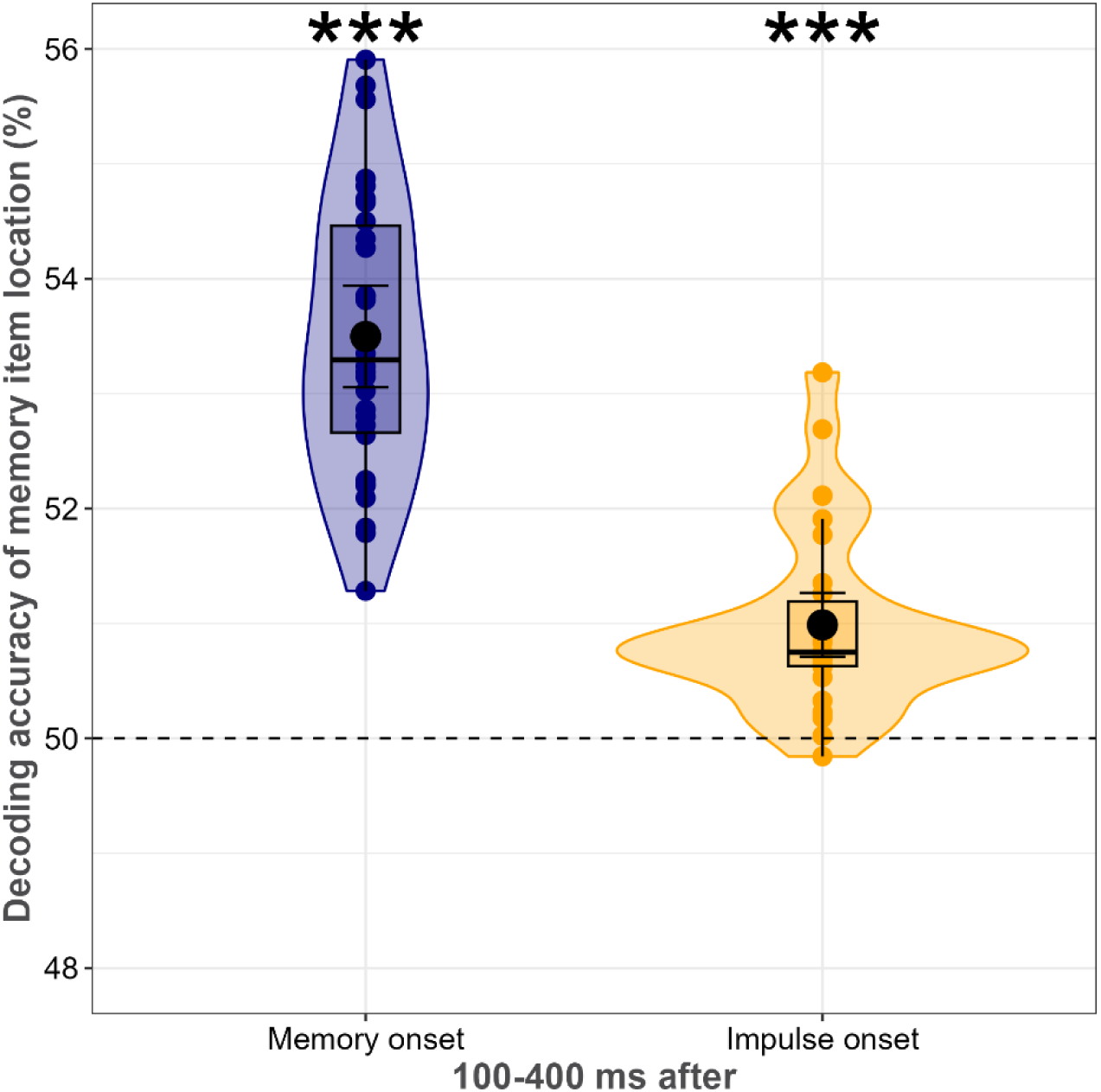
Decoding accuracy (%) of the memory item location during WM encoding and maintenance. Boxplots, violin plots, dots, error bars, and asterisks follow Figure 3.

## Discussion

By either matching or mismatching the task-irrelevant feature of spatial frequency between memory item and impulse stimulus, we aimed to manipulate the degree to which the neural impulse response is patterned by the WM-specific neural trace. Matching the spatial frequency of impulses, the task-irrelevant feature, resulted in higher decoding accuracy of the memorized orientation, the task-relevant feature. This feature-specificity of the neural impulse stimulus provides evidence that its neural response reflects the stimulus-specific neural trace in WM (Emrich et al., 2013; Pasternak & Greenlee, 2005; Postle, 2006; Stokes, 2015; Wolff et al., 2017). Notably, task-irrelevant features of the memory items were decodable during the delay period, suggesting object-based rather than feature-based WM maintenance. Furthermore, matching the spatial frequency of probes with memory items resulted in faster response times, which can be explained by a more pronounced matched-filter response, resulting in a quicker transition into the behaviorally relevant neural response that discriminates relative probe rotation (Myers et al., 2015; Sugase et al., 2008). In addition, in line with the literature (Barbosa et al., 2021; Churchland et al., 2010; Pfurtscheller et al., 1979; 1994, Min et al., 2007; Wolff et al. 2021), matching and mismatching impulse presentation equally reduced both trial-to-trial variability and alpha power most likely due to stimulus-induced neural stabilization.

The matching-dependent impulse response is clear evidence that the impulse interacts with the WM network, such that an impulse that matches this representational network better is also more effective at making its contents re-surface in the EEG. The pattern of impulse responses is thereby also in line with the idea of sensory recruitment (Emrich et al., 2013; Pasternak & Greenlee, 2005; Postle, 2006), which suggests that WM content is maintained in the sensory areas that process and represent their constituent features (e.g., line orientation in primary visual cortex). The involvement of sensory brain regions in WM is usually tested by decoding the delay activity, which can lead to mixed results (Xu, 2018). By measuring the bottom-up neural impulse response, sensory cortex can be probed without relying on measurable delay activity. Since cells responding to certain orientations are SF-specific (Daugman, 1980; Webster & de Valois, 1995), the memory items in our task should have been processed and represented by the subpopulation of neurons in visual cortex that is tuned to their specific spatial frequency. During WM maintenance, within that same population, stimulation that perturbs this network most strongly is likely to elicit the strongest memory-specific response. This is indeed what is presently found, which suggests that the orientation columns that process visual information also maintain it.

Note that the sensory recruitment explanation for why there is better decoding of matching impulses does not dictate whether the representations in WM are held in persistent activity, or in functional connectivity. In line with the former account, if orientation information is maintained by persistent activity within the same neural population during perceptual processing, perturbing these neurons could further increase their firing rate, and improve decodability. Similarly, the impulse might perturb an activity-silent functional network, which would result in the same effect. In line with the functional network account, decoding accuracy of WM content during maintenance (pre- and post-impulse) does not account for accuracy and response times (see Appendix Figure A2).

One might suppose that visual impulses may perturb neural pathways associated with sensory processing rather than WM. The matching-dependent impulse response could potentially reflect changes in synaptic efficacy within sensory pathways driven by feature-specific processing in neurons attuned to SF-specific orientations (Daugman, 1980; Webster & de Valois, 1995). However, we consider the decoding sensory processing account to be unlikely for the following reasons. A prior study by Wolff et al. (2015) implemented a single memory-item design, similar to the current study. Notably, the prior study yielded results consistent with those of a later retro-cue study (Wolff et al., 2017), where the impulse stimulus had the same SF as the memory items, similar to the “match” condition in the present study. In the retro-cue paradigm, even though impulse SF matched with relevant and no-longer relevant items, only relevant items were decodable from impulse-evoked EEG signals, indicating that matching SF does not necessarily result in perturbation of sensory history. The similarity between these results suggests that the impulse response reflects WM content rather than (sensory) presentation history in both paradigms.

Additionally, our behavioral data indicated that orientations were held in a spatial frequency-specific manner, as evidenced by faster response times when the memory spatial frequency matched that of the probe, which may be considered a hallmark of storage in WM. Furthermore, in the analysis of spatial frequency decoding, we also obtained evidence that this information was maintained in WM. Notably, in this analysis, trials were not divided between impulse-matching or mismatching, and thus, neural adaptation cannot account for the above-chance decoding of spatial frequency. Lastly, in terms of EEG signals, it would seem to us that only relatively early components are influenced by perceptual processing, occurring well before the onset of our impulse.

To further assess the possible sensory account, we also performed additional analyses, which provided evidence against the sensory account of the matching effect. First, if sensory processing were influential, higher correlations of decoding accuracy of orientations during sensory processing and after impulse presentation in the match condition would be expected due to more robust impulse perturbation in the match condition on sensory processing, but this was not observed. Correlations of decoding accuracies during sensory processing and WM maintenance were very low and did not differ between match and mismatch conditions. Second, if sensory-driven signals can account for the matching effect, better cross-generalization from stimulus-evoked to impulse-evoked signals would be expected in the match condition. The evaluation of cross-generalization showed that neural representations did not transfer from sensory processing to WM maintenance, and there was no difference between conditions. Lastly, a reanalysis of Experiment 1 from Wolff et al. (2017) showed that visually presented, but later to-be-ignored orientations were not decodable using the present dynamic baselining procedure. In line with the literature (see Wolff et al., 2020a; 2020b; Kandemir & Wolff et al., 2024), this finding suggest that the current impulse-evoked signals should similarly reflect working memories rather than sensory effects. Although these analyses cannot definitively prove that the impulse perturbs WM networks, the results provide evidence against the sensory processing account, suggesting that the impulse-matching effect is more likely attributable to WM.

Despite their behavioral irrelevance, spatial frequency and location of the memory were decodable from the impulse response further suggesting that these features were more than coincidental co-activators of the relevant orientation feature, but also a part of the memory trace itself (but see also Serences et al., 2009; Yu & Shim, 2017). This outcome is compatible with the idea that multiple features can be part of an object in memory, and that they can be encoded at minimal capacity costs (Luck & Vogel, 1997). It must be noted that in our case these multiple features shared basic, common encoding at the same location, so that the present ‘object benefits’ (i.e., the decoding of SF and location at impulse, and the reduced response times for matching probes), presumably had a stimulus-driven origin, rather than a higher-level one (cf. Balta et al., 2023). In other words, feature decoding might have been driven by their spatial integration, without necessarily requiring more conceptual object representations.

While the above-chance decoding of SF and location is in line with behavioral studies reporting maintenance (Shin & Ma, 2016; Swan et al., 2016) or attentional guidance of task-irrelevant features (Gao et al., 2016; Foerster & Schneider, 2019; but see Olivers et al., 2006), this outcome diverges from neural studies that showed little to no decoding for non-spatial task-irrelevant features during WM maintenance (Bocincova & Johnson, 2019; Yu & Shim, 2017). The divergence between our research and neural studies may have resulted from methodological differences. First, the duration of the delay period may have contributed to the above chance-level decoding of task-irrelevant features during the delay period in our study, since the neural representations of task-irrelevant features are suggested to be transient when WM load is high (Xu, 2010). Second, task-irrelevant features may not be at the center of attention during the delay period, and therefore their representations may not be maintained with sustained activity. This would make such representations difficult if not impossible to decode in fMRI and EEG without perturbation methods. Indeed, this pattern of results might suggest that irrelevant features are not maintained with sustained neural activity, and that short-term synaptic connectivity may be the mechanism. In line with this, it has been shown that the decoding of task-relevant and -irrelevant features is attenuated after WM encoding and both showed ramp-up activity before the probe onset (Bocincova & Johnson, 2019). Third, the nature of the task-irrelevant feature may explain the divergence as spatial task-irrelevant features (i.e., location) have been decoded previously during the WM delay (Foster et al., 2017; Zhou et al., 2022), while task-irrelevant color (Bocincova & Johnson, 2019) and orientation (Yu & Shim, 2017) have not been. Therefore, it could be that SF plays a special role in WM, similar to location as both SF and location determine which orientation cells process incoming perceptual information.

If the spatial frequency of the probe stimulus matched that of the memory item, behavioral performance improved, as reflected by faster response times. This can be explained by a more pronounced matched-filter response for matching stimuli, resulting in a quicker transition into the behaviorally relevant neural response that discriminates relative probe rotation (Myers et al., 2015; Sugase et al., 2008). This highlights that the WM-specificity of the impulse response reported here and previously (e.g., Fan et al., 2021; Kandemir & Akyürek, 2023; Wolff et al., 2017), should not be considered an epiphenomenon of WM-maintenance, but a useful neural mechanism for a task-specific WM-trace that is optimized for efficient readout (Myers, 2022; Nairne, 2002).

Finally, matching and mismatching impulses reduced trial-to-trial variability and alpha power, in line with the literature (Barbosa et al., 2021; Churchland et al., 2010; Pfurtscheller et al., 1979; 1994; Min et al., 2007; Wolff et al., 2021), but the matching-unspecific pattern of these effects speaks against a purely generic, SNR-mediated impulse response, as has previously been proposed (Barbosa et al., 2021). If the impulse response had not been found to be WM-specific, this would have entailed that impulse perturbation can only access representations held in persistent-activity states, much like traditional methods. However, the presently observed increased WM-content decoding of a matching impulse can only be explained by a content-specific interaction between the impulse and the current, potentially low-activity or activity-silent state of the WM network.

## Conclusion

Matching a task-irrelevant stimulus feature between the WM-item and an irrelevant impulse stimulus presented during WM maintenance increased the WM-specific neural response. Similarly, a matching probe stimulus increased WM recall as measured from a reduction in reaction times. While the impulse decreased trial-to-trial variability and alpha power, these effects were not modulated by matching between impulse and memory items. Taken together, these results suggest that the WM network maintains a stimulus-specific trace that can be targeted and readout directly via visual impulse perturbation.

## Data availability statement

Raw and processed EEG and behavioral data as well as analysis scripts will be available publicly upon publication of the manuscript on https://osf.io/7wrjh/.

## Ethics Statement

Ethical approval was obtained from the Ethical committee of psychology department at the University of Groningen prior to the data collection. All subjects signed informed consent forms prior to the data collection.

## Appendix

### Memory-Impulse SF matching effect is not driven by sensory processing

While both single and multi-item WM tasks lead to similar neural impulse responses (Wolff et al., 2015; 2017), visual impulses may interfere with sensory processing pathways. In this scenario, the impulse response dependent on SF-matching with memory orientations could result from alterations in synaptic efficacy within sensory pathways, influenced by feature-specific sensory processing in neurons that respond to particular SF and orientations (Daugman, 1980; Webster & de Valois, 1995). This hypothesis was tested by correlating decoding accuracy during encoding (100 – 400 ms after memory onset) and maintenance (100-400 ms after impulse onset), as well as through cross-generalization of stimulus-evoked and impulse-evoked neural responses.

Starting with the correlations, if stimulus-evoked sensory processing were responsible for better decoding in the match condition, greater correlations of decoding accuracy in that condition would be expected since impulse would perturb sensory-driven neural processing. However, the correlation coefficients were notably low, and there was no difference in correlations between memory-impulse spatial frequency match and mismatch conditions (*p* = .907; Figure A1a). Still, it could be argued that neural adaptation may lead to reduced firing rates in adapted networks during sensory processing, which would, theoretically, result in higher perturbation strength when a matching impulse is presented, as the matching impulse could evoke a greater change from a lower baseline of sensory-driven neural activity. First, the neural adaptation account would predict lower correlations in match condition due to a greater perturbation effect, which was not present. Second, neural adaptation should also decrease the sensory processing of impulse and evoked-impulse neural responses. It is unlikely that neural adaptation targets only sensory processing of memory orientations.

**Figure A1.**
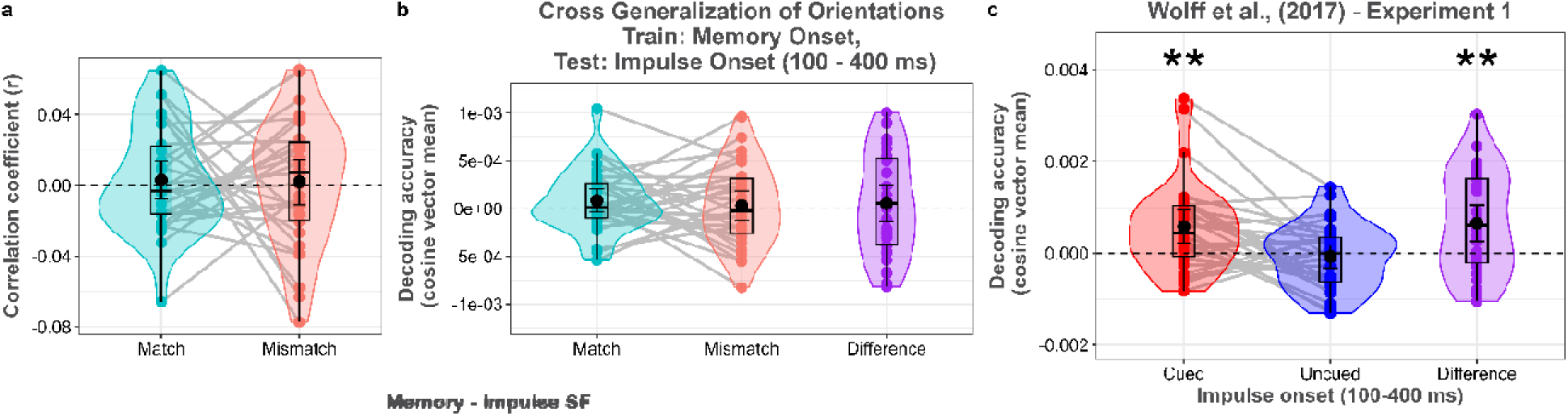
Comparison of decoding strengths of memory orientations during encoding and maintenance as a function of memory-impulse spatial frequency (SF) pairs. **a.** The correlation of decoding strengths after memory and impulse presentation. **b.** Cross-generalization of orientations from stimulus-evoked signals to impulse-evoked signals in match and mismatch conditions. None of the analyses revealed significant differences between match and mismatch conditions. **c.** Re-analysis of Wolff et al., 2017 Experiment 1 dataset using the same decoding approach as in the present study.. While cued orientations were decodable after impulse onset, decoding accuracy of uncued orientations was not significant from impulse-evoked EEG signals. Boxplots, violin plots, dots, error bars, and asterisks follow Figure 3.

Secondly, if purely sensory-driven signals could account for better decoding of memory orientations in the match condition, greater cross-generalization of memory orientations from stimulus-evoked signals (100-400 ms after memory onset) to impulse-evoked signals (100-400 ms after impulse onset) would be expected when the memory item and impulse spatial frequencies matched, compared to when they mismatched. However, the cross-generalization did not reach significance in either the match (*p* = .161; Figure A1b) or mismatch conditions (*p* = .662), and the difference was not reliable between conditions (*p* = .609). Note that this finding does not rule out the possibility that WM maintenance relies on the same neural mechanisms as sensory processing. It is possible that the evoked-impulse response reflects an interaction between the sensory processing of memory orientations and impulse processing, which may differ from pure sensory processing. However, in sum, it is implausible that the impulse-matching effect can be explained by purely sensory-driven signals.

Finally, we reanalyzed Experiment 1 from Wolff et al. (2017) using our decoding approach with dynamic baselining. If decoding accuracy for uncued items had been above chance, it would suggest evidence of decoding sensory history rather than working memory. However, our analysis revealed that while cued orientations were decodable (*p* = .004; Figure A1c), uncued orientations were not (*p* = .583). Furthermore, the difference in decoding accuracy between cued and uncued items was statistically significant (*p* = .004). These findings provide additional evidence that the main results of the current study reflect memory-related decoding rather than decoding sensory history.

### Decoding accuracy during working memory maintenance before or after impulse perturbation, as well as impulse response magnitude, does not modulate behavior

Behavioral performance was compared between high and low decoding accuracies during WM maintenance both before (0-300 ms) and after impulse onset (100-400 ms). First, the EEG signal was baselined relative to memory onset (0-200 ms), and the spatiotemporal decoding analysi was applied without mean-centering the EEG signal to maintain the stable signal during trials. A single decoding accuracy was calculated for both the pre- and post-impulse periods per participant. Following this, the decoding accuracies were z-scaled, and trials were classified into high (*z* > 1) and low (*z* < −1) categories per participant. Accuracy and response time were then compared between high and low decoding conditions for both the pre- and post-impulse periods.

Only correct and valid (*RT* > 150 ms) trials were used in response time analysis. No significant differences between conditions were revealed by permutation T-tests, suggesting that the extent to which WM content is actively retained did not modulate behavioral performance (*p*s > .05; Figure A2a&b, A2d&e). The results remained the same when the trials were split using a median-split method. Finally, the impulse response magnitude was calculated for each trial by subtracting the decoding accuracy of the pre-impulse period from that of the post-impulse period. The same strategy was then applied to generate high and low impulse response magnitude conditions. Again, no significant difference between conditions was found (*p*s > .05; Figure A2c&f).

**Figure A2.**
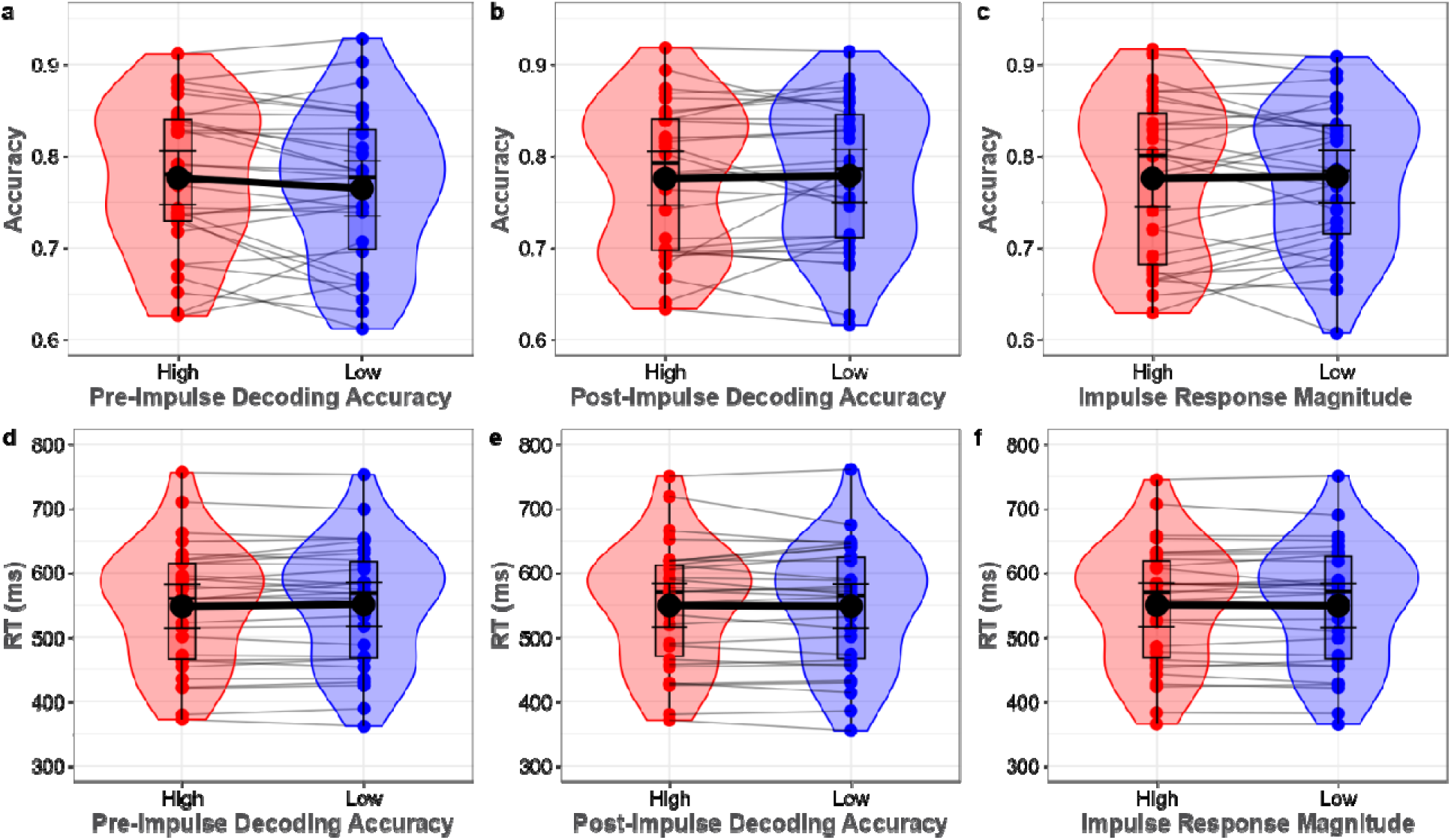
Behavioral performance between high and low decoding accuracy **a.,d.** pre-impulse and **b., e.** post-impulse period and **c., f.** the difference between post- and pre-impulse period. Boxplots, violin plots, dots, error bars, and asterisks follow Figure 3.

## Footnotes

1 Wolff et al., 2020b also showed that auditory WM representations can be perturbed by both auditory and visual impulses. This may however suggest that auditory information was visualized in their task. Furthermore, the modality-matching (i.e., auditory) impulse perturbed auditory representations better than the visual impulse, which is in line with our hypothesis.

2 By “integral,” we mean that spatial frequency and orientation are part of the same visual object in the stimulus. Although these features are technically separable, in this context they are treated as jointly perceived aspects of a single item.

